# *Begonia manicata* genome sequence reveals genetic basis underlying ornamental pigmentation

**DOI:** 10.64898/2026.02.26.708300

**Authors:** Victoria Fischer, Chiara Marie Dassow, Boas Pucker

**Author notes:** contributed equally (Victoria Fischer, Chiara Marie Dassow).

## Abstract

Plant genome sequences provide access to the gene repertoire of a species. This facilitates basic research, biotechnological processes, or horticultural applications. Here, we present the genome sequence of *Begonia manicata* and unravel the genes underlying the pigmentation of red structures emerging from its leaves and stems. Structural genes of the anthocyanin biosynthesis and corresponding regulatory genes were discovered to be upregulated in these red structures suggesting that the pigmentation is caused by the accumulation of anthocyanins. Our work provides a resource for future studies on pigmentation of Begoniaceae.

## Introduction

*Begonia* is a hyper-diverse genus comprising more than 2,000 described species and represents one of the fastest diversifying lineages of tropical plants (Hughes *et al*., 2015; Christenhusz & Byng, 2016; Moonlight *et al*., 2018). Characteristic for *Begonia* are asymmetric leaves, monoecious flowers with unequal tepals, and an outstanding morphological diversity (Permata & Susandarini, 2022), which poses a challenge for taxonomic classification without molecular and genomic data (Moonlight *et al*., 2018; Begonia Phylogeny Group *et al*., 2022). This diversity is believed to result from polyploidization events including species-specific whole genome duplications (Li *et al*., 2022; Campos-Dominguez *et al*., 2022). The genome sequence of *Begonia fimbristipula* offered additional insights into structural variations (Xiao *et al*., 2025). Due to their showy flowers, impressive floral diversity, and fascinating leaf structures, *Begonia* species are also cultivated as ornamental plants (Permata & Susandarini, 2022). In Southeast Asia, these plants are part of the traditional medicine (Borah *et al*., 2025). Both, ornamental value and medicinal relevance, can probably be attributed to the flavonoids produced by *Begonia* species. Previous studies already explored selected genes of the flavonoid biosynthesis (Emelianova *et al*., 2021).

A major group of compounds derived from the flavonoid biosynthesis are anthocyanins, responsible for the striking colors of many plant structures (Grünig *et al*., 2025). The biosynthesis of anthocyanins competes with the synthesis of other classes of flavonoids including flavones, flavonols, and proanthocyanidins (Winkel-Shirley, 2001; Grotewold, 2006; Choudhary & Pucker, 2024). The biosynthesis of anthocyanins requires a number of enzymes including CHS, CHI, F3H, DFR, ANS, arGST, and UGT (Eichenberger *et al*., 2023; Grünig *et al*., 2025). In addition, F3’H and F3’5’H can catalyze additional hydroxylations of the B ring, which shifts the color of the resulting anthocyanin from orange to pink or blue (Schwinn *et al*., 2014). After synthesis, anthocyanins can be decorated with chemical groups including glycosylation, acylation, and methylation leading to a diverse mixture of different derivatives (Grünig *et al*., 2025). After synthesis at the cytosolic side of the endoplasmatic reticulum, anthocyanins are transported into the central vacuole (Poustka *et al*., 2007; Pucker & Selmar, 2022). The structural genes of the anthocyanin biosynthesis are controlled by a complex of transcription factors (MBW complex), comprising members of the MYB, bHLH, and WD40 family (Gonzalez *et al*., 2008). While the anthocyanin biosynthesis is generally well conserved between plant species, lineage-specific or species-specific variations have been reported during the last years (Marin-Recinos & Pucker, 2024; Choudhary *et al*., 2026). The biosynthesis of anthocyanins in *Begonia* is particularly interesting, because the closely related Cucurbitaceae have been reported to almost completely lack the necessary genes for this process, which makes Begoniaceae an important outgroup for evolutionary studies of anthocyanin loss (Choudhary *et al*., 2026). Besides a large number of ecological functions of anthocyanins in reproduction and stress response (Grünig *et al*., 2025), anthocyanins also have a value in biotechnological applications as antioxidants or food colorants (Butelli *et al*., 2008; Appelhagen *et al*., 2018). Unraveling any lineage-specific elements of the anthocyanin biosynthesis in Begonia would facilitate the development of novel horticultural plants through breeding and genome editing.

An available genome sequence was considered an important property of model organisms for many years, because the access to gene sequences was crucial for molecular biology. However, recent progress in long read sequencing and plant genomics enables an almost routine generation of highly continuous plant genome sequences (Pucker *et al*., 2022; de Oliveira *et al*., 2026). Therefore, it is feasible to select plants based on economically relevant properties and sequence the genome to support research directly on the species of interest.

Here, we describe the genome sequence of *Begonia manicata* and the set of protein encoding genes. Given the horticultural importance of flower coloration, genes in the flavonoid biosynthesis are identified. Over 150 years ago, *Begonia manicata* (named *Gireoudia manicata* at the time) was already investigated due to red, soft tissues arising on the stems and leaves (Weiss, 1858) (**Fig. 1**). He hypothesized that these structures develop from hair-like epidermal outgrowths. Since the molecular basis of these fascinating structures has eluded researchers, one of the objectives of this study was to unravel the gene expression patterns in these colorful outgrowths to demonstrate the advantage of having a genome sequence.

**Fig. 1:**
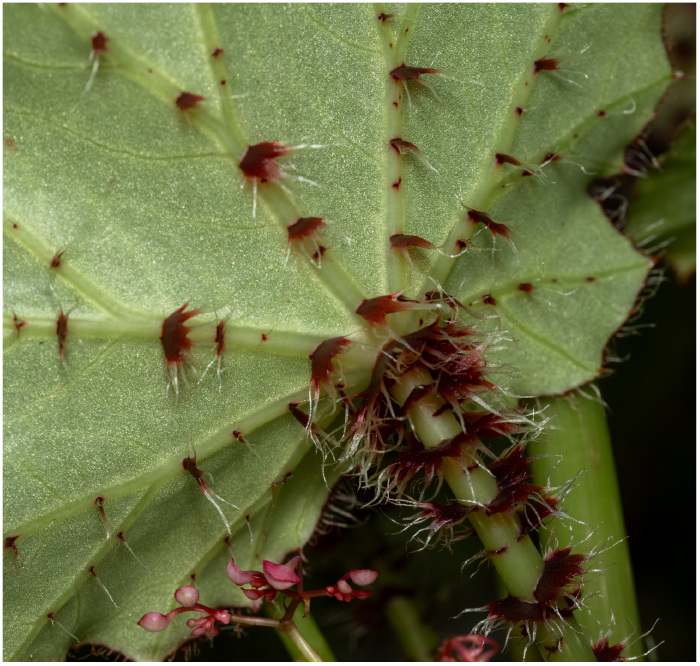
*Begonia manicata* plant (BONN-49802) with red, soft outgrowths emerging from the leaf and stems. This picture was taken in a greenhouse in the Botanical Gardens of the University of Bonn in January 2026. Photo credit: Jakob Maximilian Horz.

## Materials and Methods

### Genome sequencing

Plant material was harvested from clonally propagated plants (XX-0-BRAUN-7104585, BONN-49802) grown under greenhouse conditions (approximately 20°C, high humidity). Young leaves were selected for the extraction of high molecular weight DNA following a previously established CTAB-based protocol (Siadjeu *et al*., 2020). Quality control, library preparation, sequencing, and data analysis were conducted following a previously optimized workflow (de Oliveira *et al*., 2026). Briefly, short DNA fragments were depleted with the Short Read Eliminator kit (Pacific Biosciences). NanoDrop measurements estimated a DNA concentration, agarose gels were run to assess fragment length distribution and to rule out RNA contamination, a Qubit measurement was conducted for final DNA quantification. The library preparation with 1μg of DNA followed the SQK-LSK-114 protocol (Oxford Nanopore Technologies). Nanopore sequencing was conducted on R10 flow cells using a MinION Mk1B. Flow cells were washed and reloaded with a fresh library upon blockage of a large fraction of nanopores. Basecalling was done with dorado v0.8.3 using the model dna_r104.1_e8.2_400bps_sup@v5.0.0 and detection of modified bases (--modified-bases “5mCG_5hmCG”) on a GPU in the de.NBI cloud. Read correction with HERRO v1 was conducted (Stanojević *et al*., 2024) through dorado.

### RNA-seq and gene expression analysis

The red structures found on the leaves and the stems of several clonally propagated plants were separated by cutting them off, making sure no leaf tissue remained on the red structures. Five samples of red tissue and leaf tissue each were taken forward to RNA extraction, following the protocol in the NucloSpinRNA Plant and Fungi kit (Macherey-Nagel). For the RNA extraction of the leaves the the recommendation for *Arabidopsis thaliana* leaves (100 mg samples, 20μl PFR, 500μl PFB) was used while for the extraction of the red tissue (due to higher estimated secondary metabolite content) the recommendation for grape vine leaves (100 mg sample, 50 μl PFR, 500μl PFB) was chosen.Initial quality control was conducted on an agarose gel and Nanodrop measurement. RNA samples were sent on dry ice to a commercial service provider for quality control, cDNA synthesis, library preparation, and paired-end sequencing (Illumina NovaSeq 6000). RNA-seq data was utilized as hints for the gene prediction process and processed with kallisto v0.44 (Bray *et al*., 2016) based on identified coding sequences for quantitative analyses. Differentially expressed genes between green parts of leaves and the red structures growing on top of leaves and stems were identified with DESeq2 (Love *et al*., 2014). Correction for multiple testing was included and only genes with an adjusted p-value < 0.01 were considered for downstream analyses (Additional file H).

### Genome sequence assembly

The *Begonia manicata* genome sequence was assembled with Hifiasm v0.25.0-r726 (Cheng *et al*., 2024). HERRO-corrected reads were provided as default input (Hifi reads).The resulting assembly was assessed with contig_stats3.py (de Oliveira *et al*., 2026) and contigs shorter than 100 kb were discarded. BUSCO v6.0.0 (Tegenfeldt *et al*., 2025) was run in genome mode with eudicotyledons_odb12 as reference lineage and --skip_bbtools. Merqury v-1.3 (Rhie *et al*., 2020) was deployed to further inspect the assembly quality using an optimal k-mer size of 20.

### Structural annotation

The *Begonia manicata* genome sequence was subjected to GeMoMa v1.9 (Keilwagen *et al*., 2019) for structural annotation. Hints were provided from a range of plant species including *Argentina anserina* (GCF_933775445.1) (Christenhusz *et al*., 2023), *Malus domestica* (GCF_042453785.1) (Su *et al*., 2024), *Malus sylvestris* (GCF_916048215.2) (Ruhsam *et al*., 2022), *Rosa rugosa* (GCF_958449725.1) (Ruhsam *et al*., 2024), *Rosa chinensis* (GCF_002994745.2) (Raymond *et al*., 2018), *Prunus speciosa* (GCA_041154625.1) (Fujiwara *et al*., 2025), *Begonia darthvaderiana* (CNA0013973) (Li *et al*., 2022), *Begonia loranthoides* (CNA0013974) (Li *et al*., 2022), *Begonia masoniana* (CNA0013975) (Li *et al*., 2022), and *Begonia peltatifolia* (CNA0013976) (Li *et al*., 2022). In addition, hints derived from RNA-seq reads, that were previously mapped onto the genome assembly with HISAT2 v2.2.1, (Additional file C) were supplied and GeMoMa was run with the parameters ‘r=MAPPED and ERE.m’ using the filter GAF.f=“start==‘M’ and stop==‘*’ and (score/aa>=‘0.5’)”. Annotation completeness was assessed with BUSCO v6.0.0 (Tegenfeldt *et al*., 2025) in protein mode with eudicotyledons_odb12 as the reference. The GAF module was run to polish the annotation results with: f=“start==‘M’ and stop==‘*’ and aa>=15 and avgCov>0 and (isNaN(score) or score/aa>=‘3.25’)” atf=“tie==1 or sumWeight>1”. Finally, GeMoMa AnnotationFinalizer was utilized to rename genes. CDS and polypeptide sequences were extracted with the module GeMoMa Extractor. On the extracted protein sequences, BUSCO v6.0.0 (Tegenfeldt *et al*., 2025) was run in protein mode with eudicotyledons_odb12 as reference lineage to assess completeness.

### Functional annotation

The polypeptide sequences of the annotated Begonia manicata genome sequence were subjected to a range of tools for the assignment of functional annotation terms. A general annotation was obtained with construct_anno.py based on orthology of sequences to well studied *Arabidopsis thaliana* sequences of Araport11 (Pucker & Iorizzo, 2023). Flavonoid biosynthesis associated genes were annotated in detail using KIPEs v3 with the flavonoid bait and residue set v3.4 and default parameters (Rempel *et al*., 2023). The two transcription factor families associated with the flavonoid biosynthesis, bHLH and MYB, were annotated with the bHLH_annotator v1.04 (Thoben & Pucker, 2023) and MYB_annotator v1.0.3 (Pucker, 2022), respectively. The bHLH_annotator was run with the arguments --keepnames, --parallel and --mode_aln mafft (Additional file J). The MYB_annotator was run with --keepnames and with the *A. thaliana* reference MYB sequences provided for context (Additional file I). TTG1 ortholog candidates were identified via BLASTp v2.13.0+(Altschul *et al*., 1990, 1997) with the *A. thaliana* TTG1 polypeptide sequence (NP_197840.1). The B. manicata candidate sequences were aligned with a set of TTG1 and TTG1-like polypeptide sequences via MAFFT v7 (Katoh & Standley, 2013). The alignment was trimmed with algntrim.py (Pucker & Iorizzo, 2023) to remove almost empty columns. IQ-TREE v1.6.12 (Nguyen *et al*., 2015) was applied to infer a phylogeny based on the polypeptide sequence alignment. The model finder identified JTT+F+R6 as the best fit thus it was used for the final tree construction (Additional file E).

## Results & Discussion

### Assembly and annotation of the *Begonia manicata* genome sequence

The *Begonia manicata* genome sequence was assembled based on ONT long reads (Additional file D, (Fischer *et al*., 2026)). The performance of different assembly strategies was evaluated. The Hifiasm assembly showed overall a high level of continuity and appeared to have resolved the full *B. manicata* genome. A total of 602 Mbp were represented in 86 contigs with an N50 of 37.4 Mbp. The discovery of 98.5% complete BUSCO genes indicated a high level of completeness and Merqury supported this with a QV score of 66. Therefore, this assembly was selected as representative genome sequence for *Begonia manicata* and further analyzed in the following steps of structural and functional annotation. In total, 27,290 protein encoding genes were identified based on hints from closely related plant species and RNA-seq. The final set of predicted polypeptide sequences was analyzed with BUSCO revealing a completeness of 95.4% (C:95.4%[S:83.1%,D:12.3%],F:1.8%,M:2.8%,n:2805).

The *B. manicata* genome sequence was compared against other available *Begonia* genome sequences (Li *et al*., 2022; Xiao *et al*., 2025) (Additional file A). While the assembly size is dependent on the genome size of the respective species, continuity and completeness of an assembly can be compared across closely related species. The N50 size of the *B. manicata* assembly (37.4 Mbp) is in the same range as the best available *Begonia* genome sequences (approximately 20-50 Mbp). However, it is important to point out that the *B. manicata* assembly comprises only contiguous sequences, while the other available assemblies are the product of a scaffolding process that introduced gaps.The percentage of detected complete BUSCO genes is the highest for the *B. manicata* assembly (this study) followed by the *B. loranthoides* (Li *et al*., 2022) assembly. The availability of multiple independent genome sequences for different species in a diverse genus like Begonia is a first step towards the construction of a pangenome. Resolving the phylogenetic relationships and understanding a potential adaptive radiation in this species-rich genus will benefit from an additional genome sequence.

### Exploring the anthocyanin biosynthesis in *Begonia manicata*

Structural genes involved in the anthocyanin biosynthesis in *B. manicata* were identified with KIPEs, while transcriptional regulators were identified based on orthology to previously characterized sequences from other plant species (**Fig. 2**, Additional file B). At least one candidate was identified for each central step in the pathway. Given that *B. manicata* displays colorful red structures on leaves and stems, a functional anthocyanin biosynthesis was expected.

**Fig. 2:**
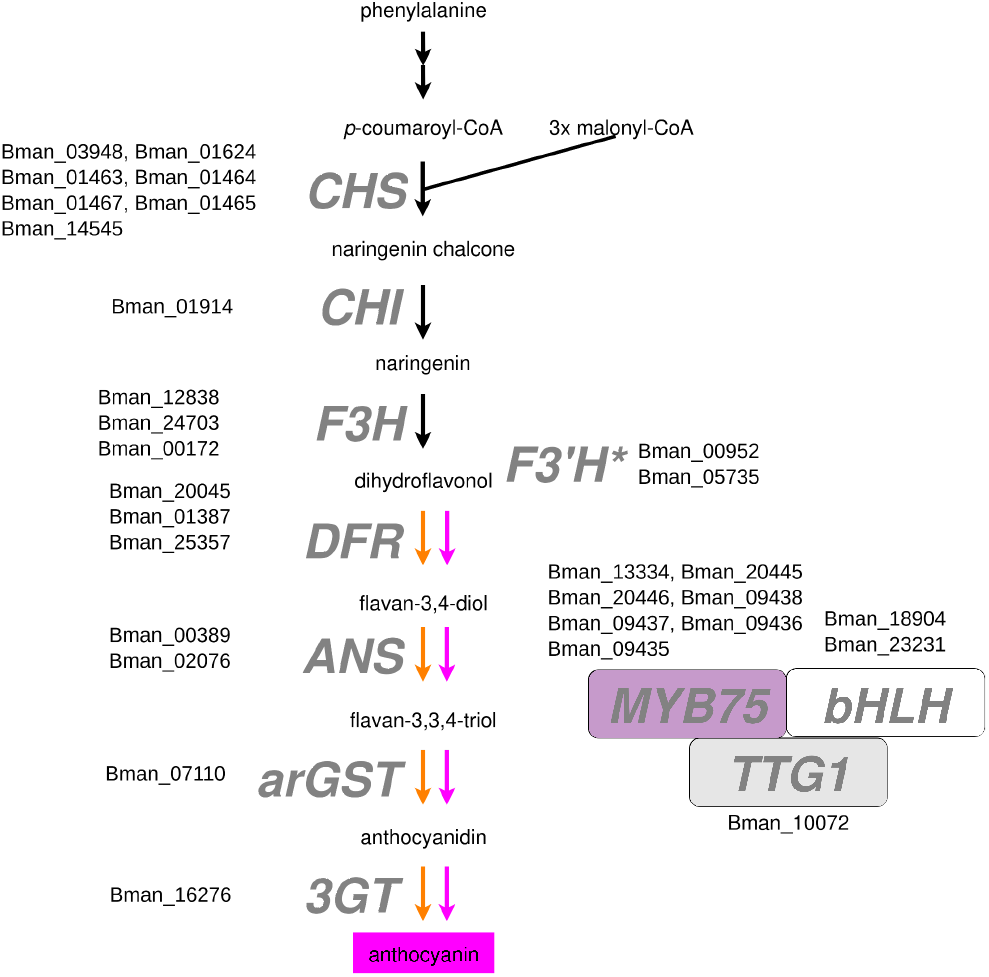
Flavonoid biosynthesis pathway in *Begonia manicata*. The candidate genes were identified with KIPEs. If amino acid residues considered functionally important are missing, the candidate is marked with an asterisk. The general figure design is inspired by (Horz *et al*., 2025). *CHS*, chalcone synthase; *CHI*, chalcone isomerase; *F3H*, flavanone 3-hydroxylase; *F3’H*, flavonoid 3’-hydroxylase; *DFR*, dihydroflavonol 3-reductase; *ANS*, anthocyanidin synthase; *arGST*, anthocyanin-related glutathione S-transferase; *3GT*, UDP-dependent anthocyanidin 3-O-glucosyltransferase; *MYB*, Myeloblastosis; *bHLH*, basic helix-loop-helix; *TTG1, TRANSPARENT TESTA GLABRA 1*.

The best F3’5’H candidate lacks crucial amino acid residues (Rempel *et al*., 2023). This is an enzyme that would be required for the biosynthesis of blue pigments. Based on the reddish pigmentation of structures on leaves and stem as well as the flower color, there is no phenotypic evidence that F3’5’H is present and active in this species.

Multiple copies for many steps in the flavonoid biosynthesis is a common pattern seen in many plant species and might even be fundamental for evolution of the specialized plant metabolism (Panchy *et al*., 2016; Rempel *et al*., 2023; Grünig *et al*., 2025). Previous reports revealed that *Begonia* species are generally showing multiple copies of genes encoding enzymes for certain steps in the flavonoid biosynthesis pathway (Emelianova *et al*., 2021; Choudhary *et al*., 2026). The chalcone synthase has been reported to show several duplications in *B. masoniana* with relaxed selection and differential expression of these paralogs (Emelianova *et al*., 2021).

### Differential gene expression between green and red leaf structures

A total of 8380 transcripts with a differential abundance were observed between the green sections of leaves and the red structures emerging from them (Additional file F). Among the differentially expressed genes are several flavonoid biosynthesis genes and their regulators including *arGST, ANS, DFR*, and an anthocyanin activating MYB. In general, structural genes in the flavonoid biosynthesis and especially those active in the anthocyanin and proanthocyanidin branches appeared substantially more active in the red structures (R1-R5) compared to green sections of the leaves (G1-G5) (**Fig. 3**, Additional file G). This aligns perfectly with the phenotype of the respective structure as a high anthocyanin content in the red structures would explain their color.

**Fig. 3:**
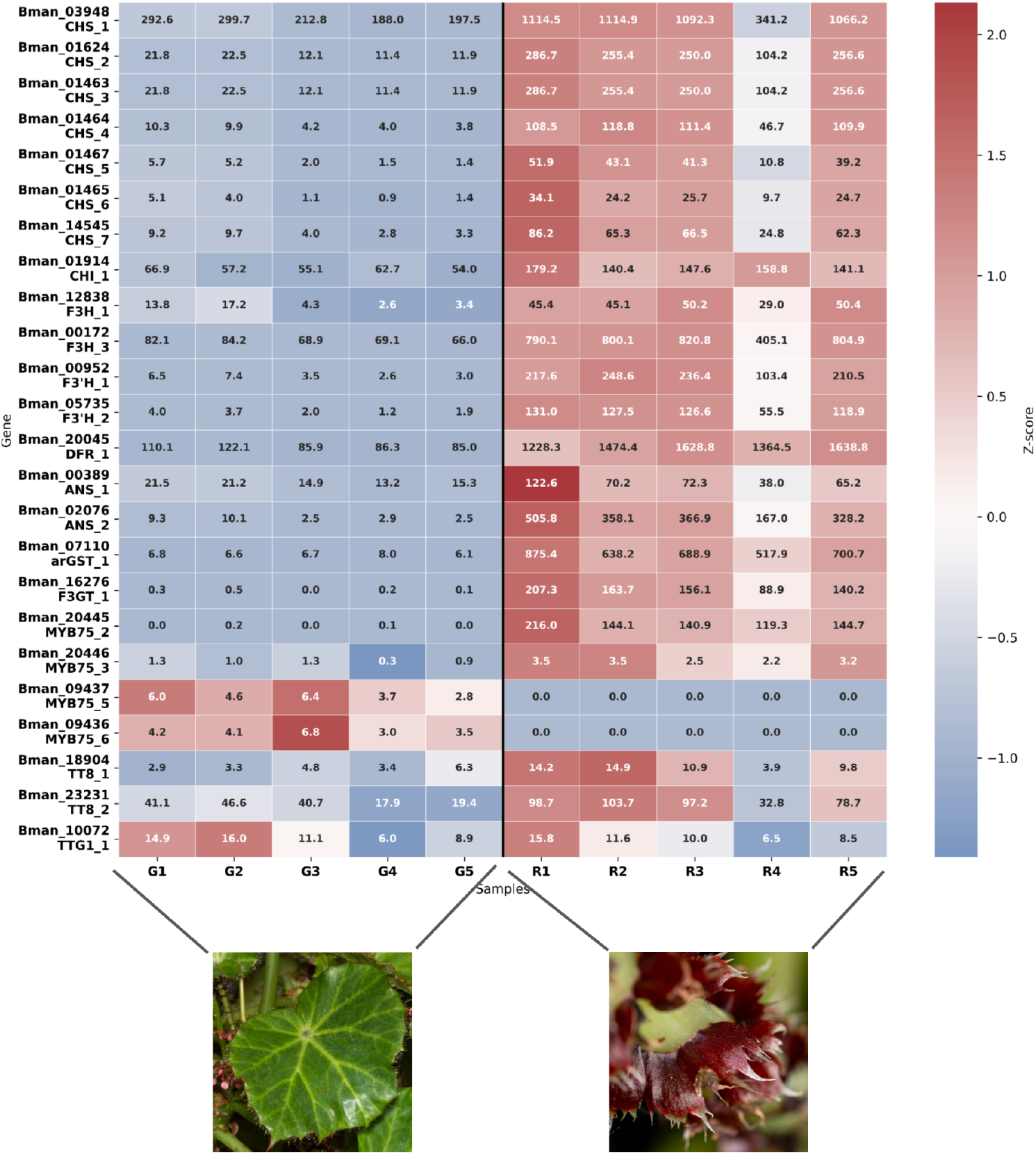
Heatmap showing the activity of flavonoid biosynthesis genes in green leaves and in red structures emerging from leaves and stems. Displayed genes are restricted to those with a combined expression of TPM > 10 across all samples. A full heatmap showing additional flavonoid biosynthesis genes is available as Additional file G. *CHS*, chalcone synthase; *CHI*, chalcone isomerase; *F3H*, flavanone 3-hydroxylase; *F3’H*, flavonoid 3’-hydroxylase; *DFR*, dihydroflavonol 3-reductase; *ANS*, anthocyanidin synthase; *arGST*, anthocyanin-related glutathione S-transferase; *3GT*, UDP-dependent anthocyanidin 3-O-glucosyltransferase; *MYB*, Myeloblastosis; *bHLH*, basic helix-loop-helix; *TTG1, TRANSPARENT TESTA 1*.

Given that there are multiple gene copies encoding enzymes for the same step in the pathway, the expression level can now reveal the relative contribution of different copies. For example, it appears that *CHS1* is substantially more active than any other *CHS* copy and that *ANS2* is substantially more active than *ANS1*. The expression patterns also allow the assignment of the transcription factors of the anthocyanin biosynthesis into different groups.

Although a complex of three transcription factors, the MBW complex, is required to activate the anthocyanin biosynthesis (Gonzalez *et al*., 2008), the individual components show different levels of specificity to the anthocyanin biosynthesis. A large number of previous studies indicate that the MYB transcription factor is the most specific component, which might not have any major functions besides activation of the anthocyanin biosynthesis genes (Marin-Recinos & Pucker, 2024). The expression pattern observed in *B. manicata* suggests that there are two groups of gene copies which differ in their tissue-specificity: MYB75_2 and MYB75_3 appear responsible for the strong activation of the anthocyanin biosynthesis in red structures emerging from leaves, while MYB75_5 and MYB75_6 appear responsible for the basal activation of the anthocyanin biosynthesis in green leaves (**Fig. 3**).

The two bHLH protein encoding *TT8* copies show slightly higher activity in the red structures with the *TT8_2* appearing as the more active copy. These expression patterns align with the expectation, because TT8 is known as an important regulator of the anthocyanin biosynthesis (Nesi *et al*., 2000; Gonzalez *et al*., 2008). Some cases of anthocyanin pigmentation loss have been attributed to TT8 (Lim *et al*., 2017; Marin-Recinos & Pucker, 2024).

The WD40 protein encoding *TTG1* shows no significantly different expression between the two groups of samples. Due to the pleiotropic roles of this transcription factor (Koornneef, 1981; Ramsay & Glover, 2005; Airoldi *et al*., 2019) a relatively constant expression across various samples is expected. For example, TTG1 has been identified as a crucial regulator of the development of trichomes and development of root hairs in *A. thaliana* (Schiefelbein *et al*., 2014; Pattanaik *et al*., 2014). It also controls the accumulation of proanthocyanidins in the seed coat of *A. thaliana* which is reflected in the seed color (Baudry *et al*., 2004; Xu *et al*., 2015). An additional function in the circadian rhythm has been reported (Airoldi *et al*., 2019).

## Conclusions

The combination of genomics and transcriptomics unraveled the genetic basis of pigmentation in *Begonia manicata*. Gene expression patterns of anthocyanin biosynthesis genes suggest that red structures emerging from green leaves are pigmented by anthocyanins. Transcription factors controlling the anthocyanin biosynthesis have been classified as tissue-specific for green sections of the leaves or the emerging red structures, respectively. The availability of sequences of these anthocyanin biosynthesis genes can facilitate future research on leaf pigmentation in *Begonia* and might support the development of horticultural innovations.

## Declarations

### Ethics approval and consent to participate

Not applicable

### Consent for publication

Not applicable

### Availability of data and materials

Sequencing data are available via ENA (PRJEB86434). Genome sequence, structural annotation, functional annotation, and expression data are available via bonndata (https://doi.org/10.60507/FK2/MOR7FZ).

### Authors’ contributions

BP designed the research project and supervised the work. CMD conducted the ONT sequencing and extracted RNA for the RNA-seq experiment. VF and CMD conducted the bioinformatic analyses. VF, CMD, and BP interpreted the results and wrote the manuscript. All authors approved the final version of the manuscript and agreed to its submission.

## Acknowledgements

This work was supported by the de.NBI Cloud within the German Network for Bioinformatics Infrastructure (de.NBI) and ELIXIR-DE (Forschungszentrum Jülich and W-de.NBI-001, W-de.NBI-004, W-de.NBI-008, W-de.NBI-010, W-de.NBI-013, W-de.NBI-014, W-de.NBI-016, W-de.NBI-022). We thank all members of the Plant Biotechnology and Bioinformatics group for their support and feedback during the process. We are grateful to Thorsten Marschall (Botanical Garden at TU Braunschweig) and the team of the University of Bonn Botanic Gardens for excellent technical support. We acknowledge support from Project DEAL and the University of Bonn for open access publication.

## References

Airoldi CA, Hearn TJ, Brockington SF, Webb AAR, Glover BJ. 2019. TTG1 proteins regulate circadian activity as well as epidermal cell fate and pigmentation. Nature Plants 5: 1145–1153. doi: 10.1038/s41477-019-0544-3.

Altschul SF, Gish W, Miller W, Myers EW, Lipman DJ. 1990. Basic local alignment search tool. Journal of Molecular Biology 215: 403–410. doi: 10.1016/S0022-2836(05)80360-2.

Altschul SF, Madden TL, Schäffer AA, Zhang J, Zhang Z, Miller W, Lipman DJ. 1997. Gapped BLAST and PSI-BLAST: a new generation of protein database search programs. Nucleic Acids Research 25: 3389–3402.

Appelhagen I, Wulff-Vester AK, Wendell M, Hvoslef-Eide A-K, Russell J, Oertel A, Martens S, Mock H-P, Martin C, Matros A. 2018. Colour bio-factories: Towards scale-up production of anthocyanins in plant cell cultures. Metabolic Engineering 48: 218–232. doi: 10.1016/j.ymben.2018.06.004.

Baudry A, Heim MA, Dubreucq B, Caboche M, Weisshaar B, Lepiniec L. 2004. TT2, TT8, and TTG1 synergistically specify the expression of BANYULS and proanthocyanidin biosynthesis in Arabidopsis thaliana. The Plant Journal: For Cell and Molecular Biology 39: 366–380. doi: 10.1111/j.1365-313X.2004.02138.x.

Begonia Phylogeny Group, Ardi W, Campos L, Chung KF, Dong W-K, Drinkwater E, Fuller D, Gagul J, Garnett G, Girmansyah D, et al. 2022. Resolving phylogenetic and taxonomic conflict in Begonia. Edinburgh Journal of Botany 79: 1–28. doi: 10.24823/ejb.2022.1928.

Borah D, Dutta R, Majumdar S, Mili C. 2025. Begonia species: a review on its ethnobotany, phytochemicals, and biological activities. Discover Plants 2: 182. doi: 10.1007/s44372-025-00272-7.

Bray NL, Pimentel H, Melsted P, Pachter L. 2016. Near-optimal probabilistic RNA-seq quantification. Nature Biotechnology 34: 525–527. doi: 10.1038/nbt.3519.

Butelli E, Titta L, Giorgio M, Mock H-P, Matros A, Peterek S, Schijlen EGWM, Hall RD, Bovy AG, Luo J, et al. 2008. Enrichment of tomato fruit with health-promoting anthocyanins by expression of select transcription factors. Nature Biotechnology 26: 1301–1308. doi: 10.1038/nbt.1506.

Campos-Dominguez L, Pellicer J, Matthews A, Leitch IJ, Kidner CA. 2022. Evolutionary patterns of genome size and chromosome number variation in Begoniaceae. Edinburgh Journal of Botany 79: 1–28. doi: 10.24823/ejb.2022.1876.

Cheng H, Asri M, Lucas J, Koren S, Li H. 2024. Scalable telomere-to-telomere assembly for diploid and polyploid genomes with double graph. Nature Methods 21: 967–970. doi: 10.1038/s41592-024-02269-8.

Choudhary N, Hagedorn M, Pucker B. 2026. Out of the blue: Family-wide loss of anthocyanin biosynthesis in Cucurbitaceae. : 2025.10.06.680802. doi: 10.1101/2025.10.06.680802.

Choudhary N, Pucker B. 2024. Conserved amino acid residues and gene expression patterns associated with the substrate preferences of the competing enzymes FLS and DFR. PLOS ONE 19: e0305837. doi: 10.1371/journal.pone.0305837.

Christenhusz MJM, Byng JW. 2016. The number of known plants species in the world and its annual increase. Phytotaxa 261: 201–217. doi: 10.11646/phytotaxa.261.3.1.

Christenhusz MJM, Leitch IJ, Royal Botanic Gardens Kew Genome Acquisition Lab, Plant Genome Sizing collective, Darwin Tree of Life Barcoding collective, Wellcome Sanger Institute Tree of Life programme, Wellcome Sanger Institute Scientific Operations: DNA Pipelines collective, Tree of Life Core Informatics collective, Darwin Tree of Life Consortium. 2023. The genome sequence of the silverweed cinquefoil, Potentilla anserina L., 1753. Wellcome Open Research 8: 464. doi: 10.12688/wellcomeopenres.19908.1.

Eichenberger M, Schwander T, Hüppi S, Kreuzer J, Mittl PRE, Peccati F, Jiménez-Osés G, Naesby M, Buller RM. 2023. The catalytic role of glutathione transferases in heterologous anthocyanin biosynthesis. Nature Catalysis 6: 927–938. doi: 10.1038/s41929-023-01018-y.

Emelianova K, Martínez Martínez A, Campos-Dominguez L, Kidner C. 2021. Multi-tissue transcriptome analysis of two Begonia species reveals dynamic patterns of evolution in the chalcone synthase gene family. Scientific Reports 11: 17773. doi: 10.1038/s41598-021-96854-y.

Fischer V, Dassow CM, Pucker B. 2026. Begonia manicata genome sequence and annotation. doi: 10.60507/FK2/MOR7FZ.

Fujiwara K, Toyoda A, Biswa BB, Kishida T, Tsuruta M, Nakamura Y, Kimura N, Kawamoto S, Sato Y, Katsuki T, et al. 2025. A Near Complete Genome Assembly of the Oshima Cherry Cerasus speciosa. Scientific Data 12: 162. doi: 10.1038/s41597-025-04388-z.

Gonzalez A, Zhao M, Leavitt JM, Lloyd AM. 2008. Regulation of the anthocyanin biosynthetic pathway by the TTG1/bHLH/Myb transcriptional complex in Arabidopsis seedlings. The Plant Journal 53: 814–827. doi: 10.1111/j.1365-313X.2007.03373.x.

Grotewold E. 2006. The genetics and biochemistry of floral pigments. Annual Review of Plant Biology 57: 761–780. doi: 10.1146/annurev.arplant.57.032905.105248.

Grünig N, Horz JM, Pucker B. 2025. Diversity and ecological functions of anthocyanins. BMC Plant Biology 26: 146. doi: 10.1186/s12870-025-08006-3.

Horz JM, Wolff K, Friedhoff R, Pucker B. 2025. Genome sequence of the ornamental plant Digitalis purpurea reveals the molecular basis of flower color and morphology variation. : 2024.02.14.580303. doi: 10.1101/2024.02.14.580303.

Hughes M, Girmansyah D, Ardi WH. 2015. Further discoveries in the ever-expanding genus Begonia (Begoniaceae): fifteen new species from Sumatra. European Journal of Taxonomy. doi: 10.5852/ejt.2015.167.

Katoh K, Standley DM. 2013. MAFFT Multiple Sequence Alignment Software Version 7: Improvements in Performance and Usability. Molecular Biology and Evolution 30: 772–780. doi: 10.1093/molbev/mst010.

Keilwagen J, Hartung F, Grau J. 2019. GeMoMa: Homology-Based Gene Prediction Utilizing Intron Position Conservation and RNA-seq Data. Methods in Molecular Biology (Clifton, N.J.) 1962: 161–177. doi: 10.1007/978-1-4939-9173-0_9.

Koornneef M. 1981. The complex syndrome of ttg mutants. Arabidopsis Information Service 18: 45–51.

Li L, Chen X, Fang D, Dong S, Guo X, Li N, Campos-Dominguez L, Wang W, Liu Y, Lang X, et al. 2022. Genomes shed light on the evolution of Begonia, a mega-diverse genus. New Phytologist 234: 295–310. doi: 10.1111/nph.17949.

Lim S-H, Kim D-H, Kim JK, Lee J-Y, Ha S-H. 2017. A Radish Basic Helix-Loop-Helix Transcription Factor, RsTT8 Acts a Positive Regulator for Anthocyanin Biosynthesis. Frontiers in Plant Science 8. doi: 10.3389/fpls.2017.01917.

Love MI, Huber W, Anders S. 2014. Moderated estimation of fold change and dispersion for RNA-seq data with DESeq2. Genome Biology 15: 550. doi: 10.1186/s13059-014-0550-8.

Marin-Recinos MF, Pucker B. 2024. Genetic factors explaining anthocyanin pigmentation differences. BMC Plant Biology 24: 627. doi: 10.1186/s12870-024-05316-w.

Moonlight PW, Ardi WH, Padilla LA, Chung K-F, Fuller D, Girmansyah D, Hollands R, Jara-Muñoz A, Kiew R, Leong W-C, et al. 2018. Dividing and conquering the fastest–growing genus: Towards a natural sectional classification of the mega–diverse genus Begonia (Begoniaceae). TAXON 67: 267–323. doi: 10.12705/672.3.

Nesi N, Debeaujon I, Jond C, Pelletier G, Caboche M, Lepiniec L. 2000. The TT8 Gene Encodes a Basic Helix-Loop-Helix Domain Protein Required for Expression of DFR and BAN Genes in Arabidopsis Siliques. The Plant Cell 12: 1863–1878.

Nguyen L-T, Schmidt HA, von Haeseler A, Minh BQ. 2015. IQ-TREE: A Fast and Effective Stochastic Algorithm for Estimating Maximum-Likelihood Phylogenies. Molecular Biology and Evolution 32: 268–274. doi: 10.1093/molbev/msu300.

de Oliveira JAVS, Choudhary N, Meckoni SN, Nowak MS, Hagedorn M, Pucker B. 2026. Cookbook for plant genome sequences. BMC Genomics. doi: 10.1186/s12864-026-12623-z.

Panchy N, Lehti-Shiu M, Shiu S-H. 2016. Evolution of Gene Duplication in Plants. Plant Physiology 171: 2294–2316. doi: 10.1104/pp.16.00523.

Pattanaik S, Patra B, Singh SK, Yuan L. 2014. An overview of the gene regulatory network controlling trichome development in the model plant, Arabidopsis. Frontiers in Plant Science 5. doi: 10.3389/fpls.2014.00259.

Permata DA, Susandarini R. 2022. Morphological diversity and phenetic relationship of wild and cultivated Begonia based on morphology and leaf venation. Biodiversitas Journal of Biological Diversity 23. doi: 10.13057/biodiv/d230235.

Poustka F, Irani NG, Feller A, Lu Y, Pourcel L, Frame K, Grotewold E. 2007. A Trafficking Pathway for Anthocyanins Overlaps with the Endoplasmic Reticulum-to-Vacuole Protein-Sorting Route in Arabidopsis and Contributes to the Formation of Vacuolar Inclusions. Plant Physiology 145: 1323–1335. doi: 10.1104/pp.107.105064.

Pucker B. 2022. Automatic identification and annotation of MYB gene family members in plants. BMC Genomics 23: 220. doi: 10.1186/s12864-022-08452-5.

Pucker B, Iorizzo M. 2023. Apiaceae FNS I originated from F3H through tandem gene duplication. PLOS ONE 18: e0280155. doi: 10.1371/journal.pone.0280155.

Pucker B, Irisarri I, Vries J de, Xu B. 2022. Plant genome sequence assembly in the era of long reads: Progress, challenges and future directions. Quantitative Plant Biology 3: e5. doi: 10.1017/qpb.2021.18.

Pucker B, Selmar D. 2022. Biochemistry and Molecular Basis of Intracellular Flavonoid Transport in Plants. 11: 963. doi: 10.3390/plants11070963.

Ramsay NA, Glover BJ. 2005. MYB-bHLH-WD40 protein complex and the evolution of cellular diversity. Trends in Plant Science 10: 63–70. doi: 10.1016/j.tplants.2004.12.011.

Raymond O, Gouzy J, Just J, Badouin H, Verdenaud M, Lemainque A, Vergne P, Moja S, Choisne N, Pont C, et al. 2018. The Rosa genome provides new insights into the domestication of modern roses. Nature Genetics 50: 772–777. doi: 10.1038/s41588-018-0110-3.

Rempel A, Choudhary N, Pucker B. 2023. KIPEs3: Automatic annotation of biosynthesis pathways. PLOS ONE 18: e0294342. doi: 10.1371/journal.pone.0294342.

Rhie A, Walenz BP, Koren S, Phillippy AM. 2020. Merqury: reference-free quality, completeness, and phasing assessment for genome assemblies. Genome Biology 21: 245. doi: 10.1186/s13059-020-02134-9.

Ruhsam M, Bell D, Hart M, Hollingsworth P. 2022. The genome sequence of the European crab apple, Malus sylvestris (L.) Mill., 1768. Wellcome Open Research 7: 296. doi: 10.12688/wellcomeopenres.18645.1.

Ruhsam M, Hollingsworth P, Royal Botanic Garden Edinburgh Genome Acquisition Lab, Plant Genome Sizing collective, Darwin Tree of Life Barcoding collective, Wellcome Sanger Institute Tree of Life Management, Samples and Laboratory team, Wellcome Sanger Institute Scientific Operations: Sequencing Operations, Wellcome Sanger Institute Tree of Life Core Informatics team, Tree of Life Core Informatics collective, et al. 2024. The genome sequence of the Japanese rose, Rosa rugosa Thunb., 1784 (Rosaceae). Wellcome Open Research. doi: 10.12688/wellcomeopenres.22910.1.

Schiefelbein J, Zheng X, Huang L. 2014. Regulation of epidermal cell fate in Arabidopsis roots: the importance of multiple feedback loops. Frontiers in Plant Science 5. doi: 10.3389/fpls.2014.00047.

Schwinn K, Miosic S, Davies K, Thill J, Gotame TP, Stich K, Halbwirth H. 2014. The B-ring hydroxylation pattern of anthocyanins can be determined through activity of the flavonoid 3′-hydroxylase on leucoanthocyanidins. Planta 240: 1003–1010. doi: 10.1007/s00425-014-2166-3.

Siadjeu C, Pucker B, Viehöver P, Albach DC, Weisshaar B. 2020. High Contiguity de novo Genome Sequence Assembly of Trifoliate Yam (Dioscorea dumetorum) Using Long Read Sequencing. Genes 11: 274. doi: 10.3390/genes11030274.

Stanojević D, Lin D, Nurk S, Sessions PF de, Šikić M. 2024. Telomere-to-Telomere Phased Genome Assembly Using HERRO-Corrected Simplex Nanopore Reads. : 2024.05.18.594796. doi: 10.1101/2024.05.18.594796.

Su Y, Yang X, Wang Y, Li J, Long Q, Cao S, Wang X, Liu Z, Huang S, Chen Z, et al. 2024. Phased telomere-to-telomere reference genome and pangenome reveal an expansion of resistance genes during apple domestication. Plant Physiology 195: 2799–2814. doi: 10.1093/plphys/kiae258.

Tegenfeldt F, Kuznetsov D, Manni M, Berkeley M, Zdobnov EM, Kriventseva EV. 2025. OrthoDB and BUSCO update: annotation of orthologs with wider sampling of genomes. Nucleic Acids Research 53: D516–D522. doi: 10.1093/nar/gkae987.

Thoben C, Pucker B. 2023. Automatic annotation of the bHLH gene family in plants. BMC Genomics 24: 780. doi: 10.1186/s12864-023-09877-2.

Weiss GA. 1858. Ueber die Entwickelungsgeschichte und den anatomi-schen Bau der handformigen Auswüchse an den Blättern und Stengeln von Gireoudia manicata Klotzsch. Acta ZooBot Austria.

Winkel-Shirley B. 2001. Flavonoid Biosynthesis. A Colorful Model for Genetics, Biochemistry, Cell Biology, and Biotechnology. Plant Physiology 126: 485–493. doi: 10.1104/pp.126.2.485.

Xiao T-W, Wang Z-F, Yan H-F. 2025. A chromosomal-level genome assembly of Begonia fimbristipula (Begoniaceae). Scientific Data 12: 429. doi: 10.1038/s41597-025-04768-5.

Xu W, Dubos C, Lepiniec L. 2015. Transcriptional control of flavonoid biosynthesis by MYB–bHLH–WDR complexes. Trends in Plant Science 20: 176–185. doi: 10.1016/j.tplants.2014.12.001.

