## Supplementary figures and images for "*Begonia manicata* genome sequence reveals genetic basis underlying ornamental pigmentation"

### Additional file E

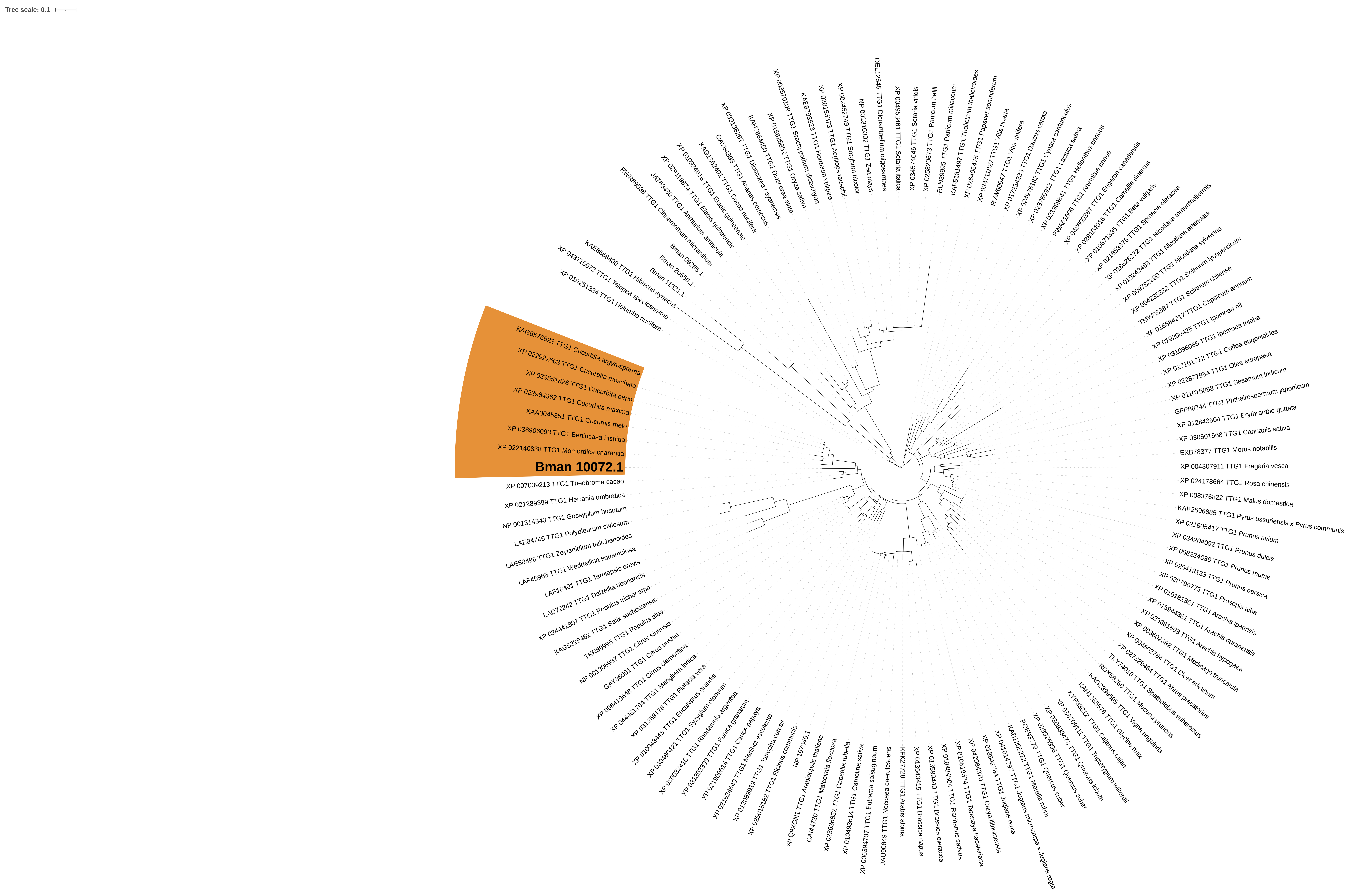

### Additional file G

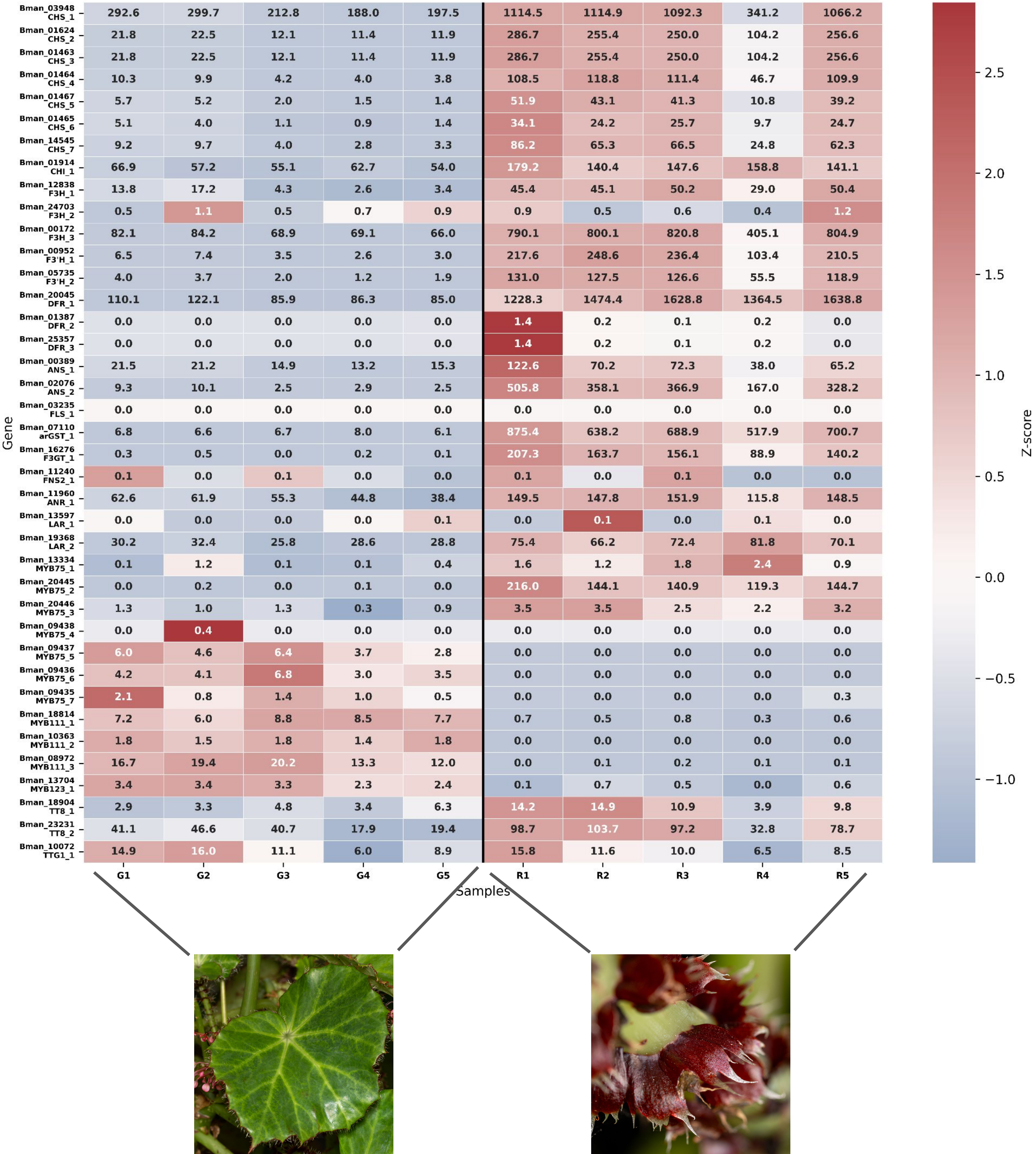

### Additional file I

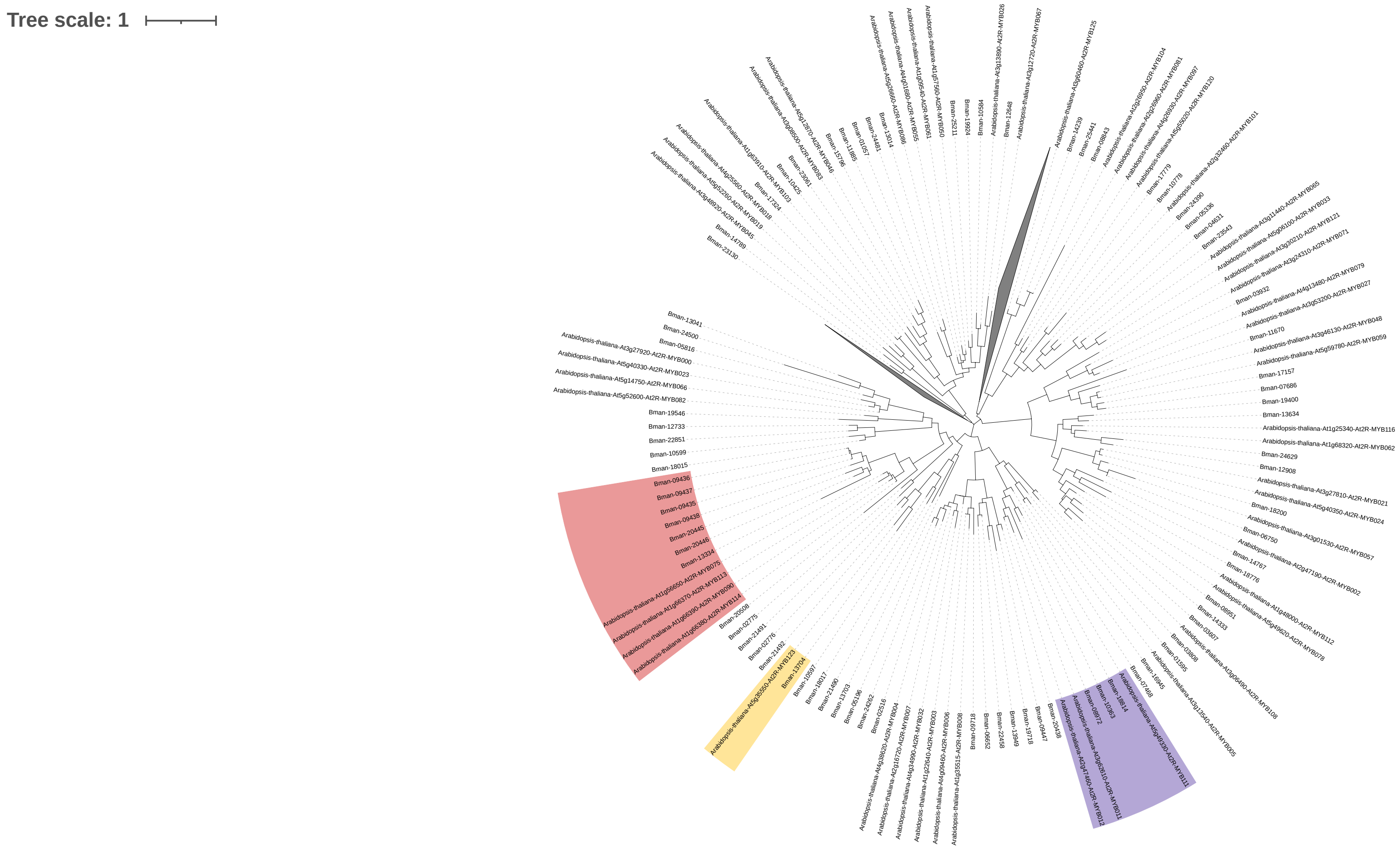

### Additional file J

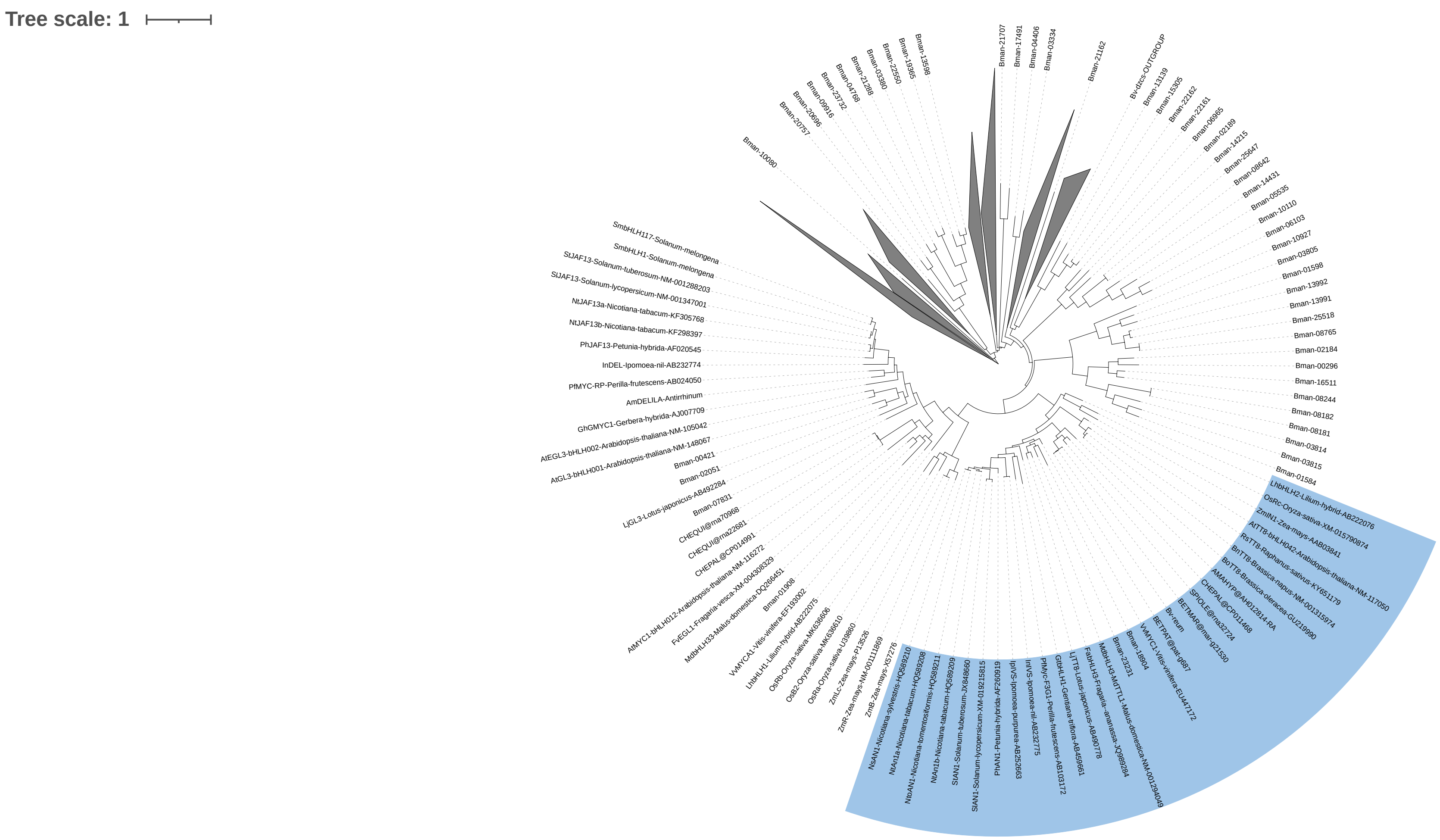
